# A synthetic integral feedback controller for robust tunable regulation in bacteria

**DOI:** 10.1101/170951

**Authors:** Gabriele Lillacci, Stephanie Aoki, David Schweingruber, Mustafa Khammash

**Affiliations:** ETH ZürichDepartment of Biosystems Science and Engineering Mattenstrasse 26, 4058 Basel, Switzerland

## Abstract

We report on the first engineered integral feedback control system in a living cell. The controller is based on the recently published antithetic integral feedback motif [1] which has been analytically shown to have favorable regulation properties. It is implemented along with test circuitry in *Escherichia coli* using seven genes and three small-molecule inducers. The closed-loop system is highly tunable, allowing a regulated protein of interest to be driven to a desired level and maintained there with precision. Realized using a sigma/anti-sigma protein pair, the integral controller ensures that regulation is maintained in the face of perturbations that lead to the regulated protein’s degradation, thus serving as a proof-of-concept prototype of integral feedback implementation in living cells. When suitably optimized, this integral controller may be utilized as a general-purpose robust regulator for genetic circuits with unknown or partially-known topologies and parameters.

## Introduction

Feedback regulation is a running theme in all of biology. The palette of regulatory strategies that employ feedback is exquisitely diverse, and can be found at every level of biological organization, from the molecular to the organismal and ecological. Among these are strategies that give rise to *homeostasis*, that defining property of organisms in which a physiological variable is maintained constant through active regulation. Maintaining homeostasis often requires that perfect or near-perfect adaptation is achieved. This means that the physiological variable adapts to a pre-established baseline after a persistent disturbance or a stimulus is encountered. The regulatory strategy accomplishes this by establishing a new balance of activities within the system so as to maintain that baseline in the presence of the disturbance or stimulus. Achieving this level of robustness requires that the regulatory strategy employs a structural motif that implements *integral feedback*. This type of feedback is dynamic and functions by measuring (directly or indirectly) a quantity that reflects the deviation from the desired baseline (setpoint), integrating that quantity over time, and then using the outcome to drive processes that counteract the deviation and drive it to zero (Fig. 1a). Integral feedback has been discovered in various biological systems including blood calcium levels in mammals [2], yeast osmoregulation [3] and bacterial chemotaxis [4].

**Figure 1:**
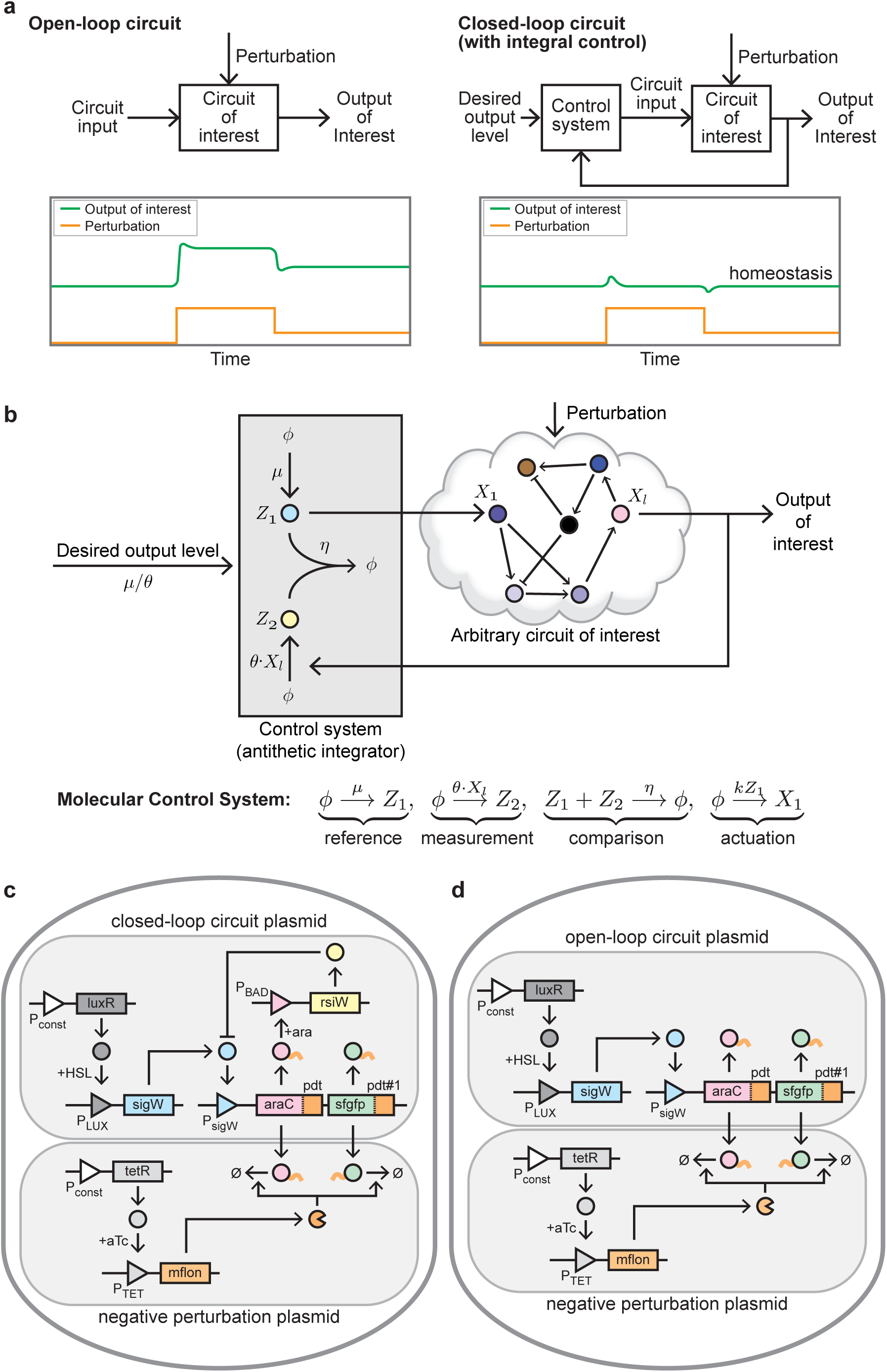
Circuit details. (a) Robust regulation of gene expression through integral feedback. The integral feedback control system computes the mathematical integral of the error between the output protein amount and the desired level, and uses the result to robustly regulate the output protein of a circuit of interest. The closed-loop system can keep the output level at the desired level in the face of external perturbations, while an open-loop circuit lacking integral feedback can not. (b) Antithetic integral feedback topology. Two controller proteins with the property of inhibiting each other’s activity (“annihilation”) are used to robustly regulate an arbitrary circuit of interest. This regulation strategy was proposed in [1] and shown to have favorable properties. (c) Closed-loop antithetic integral feedback circuit and (d) Open-loop control circuit. Triangles denote promoters, squares are genes and circles represent proteins. Pointed arrows signify activation, flat arrows inhibition. The large light grey boxes enclose the parts on the same plasmid.

While endogenous feedback strategies in biology have been shaped by nature’s evolutionary explorations over several billions years, it is only recently that genetic engineering methods are enabling their *de novo* engineering in living cells. It is now feasible to design and implement new feedback control systems in living cells which attain precise, robust, and high performance steering of cellular processes, with potential for many novel applications in Biotechnology. We refer to the technology to build such control systems as *Cybergenetics* [1]—a genetics era realization of Norbert Wiener’s Cybernetics vision [5]. Over the last few years, several synthetic feedback circuits have been engineered (see [6] for a review). However, up to now the synthetic implementation of integral feedback circuits in living cells has remained elusive, despite its widely recognized benefits and potential applications [7]. In [8, 9, 10] mechanisms for robust perfect adaptation and integral feedback were proposed and theoretically investigated. In [1], we introduced a new motif for such feedback called *antithetic integral control* and theoretically demonstrated its favorable robustness properties. We showed that such properties are maintained—and even enhanced—in the stochastic regime.

Here, we present a synthetic bacterial gene expression controller that realizes integral feedback in *Escherichia coli* based on the antithetic motif in [1]. A combinatorial approach of theoretical analysis and synthetic biology was used to realize this circuit. The controller is tunable using small-molecule inducers and can ensure than an output protein of interest is kept at a constant level in the face of perturbations that lead to its degradation. Our device should enable robust regulation of a desired genetic circuit in live bacteria.

## Results

### Circuit

The *antithetic* design [1] requires two controller proteins that “annihilate” or abolish each other’s biological activity upon reacting, for example by forming an inert dimer (Fig. 1b). To achieve this, we used a previously reported pair of *Bacillus subtilis σ* and anti-*σ* factors, *sigW* and *rsiW*, which has been shown to display annihilation *in vivo* [11]. Promoter P_*sigW*_ that responds to SigW drives the controlled gene, a bicistronic construct comprising the *E. coli* transcription factor *araC* and *sfgfp* (superfolder green fluorescent protein) [12]. *araC* implements the feedback, as AraC abundance dictates the activity of its cognate P_BAD_ promoter, which in turn drives *rsiW* expression. SfGFP is the regulated protein of interest, and in this configuration its amount should be proportional to that of AraC. The response of P_BAD_ to AraC can be adjusted with arabinose (ARA): higher ARA concentrations correspond to higher P_BAD_ activity for a given AraC concentration. The expression of *sigW* is controlled by the N-(3-oxohexanoyl)-L-homoserine lactone (HSL) inducible *Vibrio fisheri* quorum sensing pathway promoter P_LUX_ [13]. Higher concentrations of HSL result in higher P_LUX_ activity and increased production of SigW. To test the system’s response to a pertubation, we use the orthogonal *Mesoplasma florum* protease *mf*-Lon [14] driven by an anhydrotetracycline (aTc) inducible promoter (P_TET_). The *mf*-Lon protease recognizes cognate Pdt degradation tags [14], copies of which have been appended to the C-termini of AraC and sfGFP. As a result, when the protease is induced, the degradation rate of AraC and sfGFP is increased. We will refer to this controller as the *closed-loop* circuit (Fig. 1c). For comparison, we also constructed an *open-loop* circuit, in which the feedback has been disabled by removing P_BAD_-*rsiW* (Fig. 1d).

### Tunability

Next, we sought to identify the range of inducer concentrations in which the output protein in the closed-loop circuit could be regulated. For our purposes it was also important to select set points with gene expression levels high enough for detection of sfGFP in single cells by flow cytometry but low enough as to avoid an appreciable fitness burden. For all experiments, we diluted overnight stationary-phase cultures into induction media and grew them to exponential-phase with dilutions at regular intervals to maintain continuous low-density growth (Supplementary Section S2). HSL and ARA were used to independently and systematically tune expression levels of *sigW* and *rsiW*, respectively, in all combinations (Fig. 2a). As expected, increasing SigW production corresponded to an increase in sfGFP fluorescence, whereas increasing RsiW production corresponded to a decrease. These results showed that the circuit is usable over a wide range of conditions, consistent with recent work by Annunziata *et al.* showing output tunability in response to different ratios of sigma and anti-sigma from *Pseudomonas fluorescens Pf-5* in an open-loop circuit [15]. The data also suggested that the range of HSL concentrations in which the set point varies approximately linearly with HSL lies approximately between 0 and 20 nM for the tested ARA concentrations. At higher HSL concentrations the increase becomes incrementally smaller. We conjecture that this is due to saturation of the output P_*sigW*_ promoter. Away from saturation, for a fixed ARA concentration the set point depends linearly on HSL concentration (Fig. 2b, left panel). The slope of this linear dependence is inversely proportional to the ARA concentration (Fig. 2b, right panel). This shows that the set point is proportional to the ratio of the two inducers, in perfect agreement with the theory in [1].

**Figure 2:**
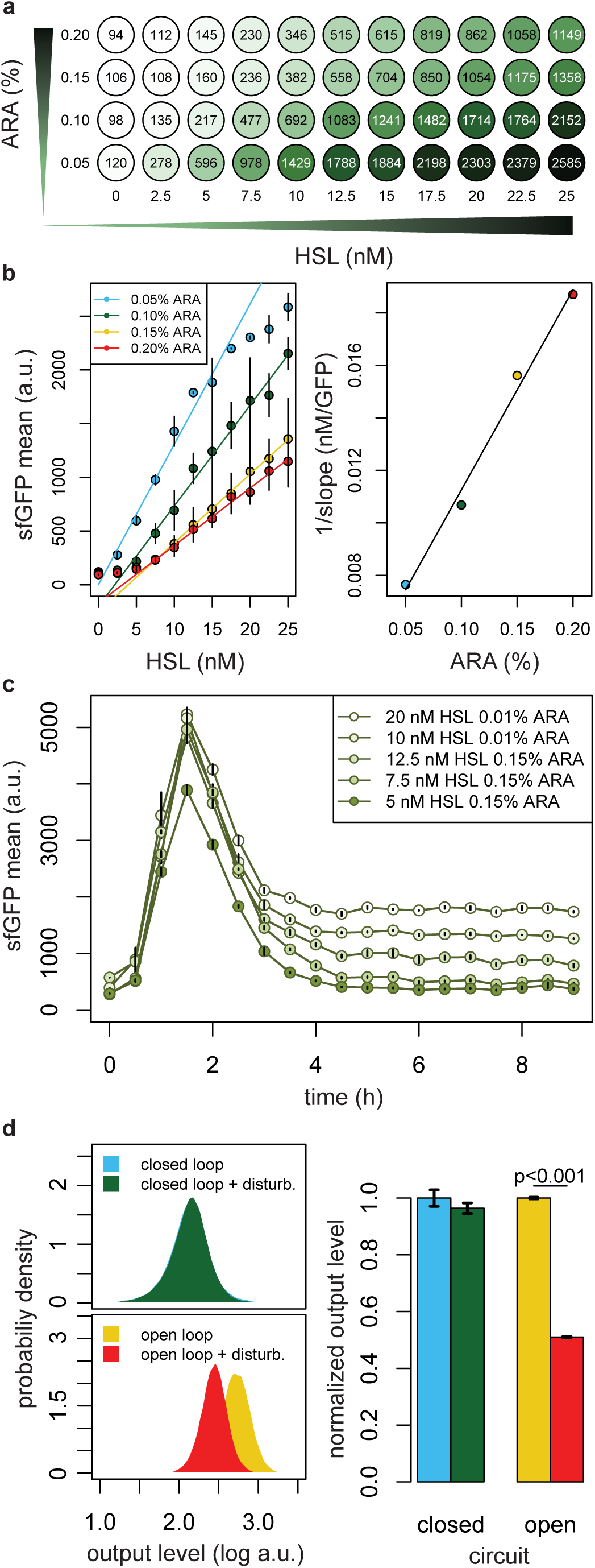
Data. (a) Response of closed-loop circuit to HSL and ARA. Means of steady-state sfGFP fluorescence distributions for the corresponding concentrations of HSL and ARA (average of 2 replicates). (b) Output steady state dependency on HSL and ARA. Left panel: means of closed-loop circuit steady-state sfGFP fluorescence distributions as a function of HSL concentration, for different ARA concentrations. Right panel: inverse of the slope of dependency between output and HSL as a function of ARA concentration. (c) Dynamic response of closed-loop circuit to HSL induction. Means of sfGFP fluorescence distributions as a function of time. Error bars denote standard deviation (*N* = 3). (d) Output steady states of closed- and open-loop circuits in presence of protease perturbation. Left panel: steady-state distributions of sfGFP fluorescence. Right panel: average fluorescence level normalized to the pre-disturbance level for each circuit; error bars denote standard deviation (*N* = 3).

### Set point tracking

We further tested the ability of the closed-loop circuit to maintain a set point by performing a series of dynamic experiments over a range of set points identified in Fig. 2a. HSL was added to cells grown in ARA and expression of sfGFP was followed over time (Fig. 2c). Fluorescence intensities were detectable over background and unimodal (Supplementary Fig. S10). The data shows that it is possible to finely and reproducibly control the steady-state levels of sfGFP and to maintain such levels over long periods of time. A transient increase in reporter fluorescence upon HSL induction was observed before cells reached steady state expression levels. However, this increase in sfGFP fluorescence is likely a result of the rapid shift from stationary-phase to exponential-phase, as the large peak is not observed in cultures pregrown to exponential phase before the application of the induction (Supplementary Fig. S9).

### Disturbance rejection

To show the effect of integral feedback, we sought to demonstrate settings for which perfect or near-perfect adaptation is achieved in the presence of a *persistent* disturbance. We measured the steady-state sfGFP fluorescence with and without aTc-induction of the *mf*-Lon protease, in both the closed-loop and the open-loop circuits. To quantify the improvement in disturbance rejection performance, we computed the difference between the normalized post-disturbance output decrease in the open-loop and in the closed-loop. The closed-loop circuit showed no change in sfGFP fluorescence after aTc-induction of the protease, while the open-loop circuit showed a 50% decrease (Fig. 2d). Our results suggest that the closed-loop circuit responds to the loss of AraC by decreasing production of anti-*σ* factor RsiW leading to an increase in active SigW and the recovery of AraC to pre-perturbation levels. In contrast, the open-loop circuit lacks the feedback regulation and is unable to respond to the perturbation.

## Discussion

We have engineered and tested a genetic circuit in *E. coli* that implements integral feedback. The circuit controls a gene of interest by driving its output precisely and robustly to a desired set point and clamping it at that value over time. We have demonstrated that the circuit is capable of perfect adaptation when the output protein of interest is subjected to a large and persistent disturbance in the form of a protease that degrades it.

The desired set point is achieved by adjusting any one of two small molecule inducers, with the ratio of their concentration defining the value of set point. This fact has significant implications when the copy number of the plasmid encoding the control circuit varies considerably between cells, as would be the case in mammalian transient transfections. In this case, the activity of the two genes whose promoter responds to the two inducers will vary commensurately, and so long as they are implemented on the same plasmid, the ratio of their response will remain unchanged, making the set point immune to plasmid copy number variability.

While the feedback circuit achieves disturbance rejection that is superior to the open-loop circuit (no feedback), our analysis suggests that the regime over which perfect or near perfect rejection is observed depends on several factors, including the size of the disturbance relative to the set point, the saturation of the genes that carry out the sensing and actuation, and the dilution rate due to cell growth. Optimizing the circuit by increasing the dynamic range of sensor and actuator genes (*sigW* and *rsiW*) will simultaneously ensure a larger dynamic range of operation and better disturbance rejection (including perfect adaptation) of larger disturbances.

The effect of dilution can also be counteracted by suitably choosing the controller parameters without affecting the set point. The original theoretical analysis in [1] showed that in the ideal case when the controller proteins are lost only due to annihilation, the set point is determined by the ratio of two parameters *µ/θ*, and that perfect adaptation will be achieved regardless of the value of the other parameters *η* and *k* (assuming no saturation). When dilution is present, however, all controller parameters impact the disturbance rejection performance, and by selecting these parameters in a suitable region of the parmeter space, perfect adaptation can be restored without altering the set point. This can be achieved in several different ways without requiring additional circuitry (Supplementary Text S1.1.1). For example if we choose *θk* sufficiently large while either keeping *η* sufficiently large or *k* sufficiently small, the impact of dilution on deteriorating disturbance rejection can be made arbitrarily small. Other approaches that require additional circuitry have been proposed in [16]. In Supplementary Text S1, we further show that by appropriately tuning the controller using practically realizable parameter values, one can circumvent the loss of perfect adaptation due to dilution even for fast growing cells.

The ability to engineer molecular homeostasis through integral feedback provides a powerful new tool in the synthetic biology arsenal, and should find wide application in all scenarios in which protein expression must remain tightly regulated at the desired level, independent of other intervening processes. In metabolic engineering, for instance, robust set point regulation of key enzymes could be used to redirect metabolic fluxes to maximize yield and minimize host toxicity. Probing endogenous pathways could also benefit from such regulation, because compensatory cellular mechanisms tend to alter expression in complex ways. A mammalian analog of our synthetic integral feedback module is currently under development. Such a circuit could also offer exciting perspectives for molecular therapy in all those conditions that result from disruption of homeostatic regulation by enabling the implementation of a fully autonomous and personalized intervention.

Beyond utility, the engineering of an integral feedback controller in single living cells is an important and natural milestone in the path to build sophisticated dynamic control systems in living systems. But arriving at a functioning proof-of-concept integral circuit was challenging both conceptually and experimentally, necessitating the development of a new control motif and a suitable control theory adapted to it. Moving beyond this first step towards more advanced cybergenetic controllers will require a better understanding of the unique environment of the living cell, together with powerful theories and experimental tools to support rational design within this environment. The challenges abound, but so do the promises.

## Acknowledgments

We thank Timothy Frei for assistance in building some the biological parts used in the final circuit. We thank Dr. Daniel Meyer for assistance with the programming of the robotic platform for the high-throughput assays.

